# Sub-voxel light-sheet microscopy for high-resolution, high-throughput volumetric imaging of large biomedical specimens

**DOI:** 10.1101/255695

**Authors:** Peng Fei, Jun Nie, Juhyun Lee, Yichen Ding, Shuoran Li, Hao Zhang, Masaya Hagiwara, Tingting Yu, Tatiana Segura, Chih-Ming Ho, Dan Zhu, Tzung K. Hsiai

**Affiliations:** School of Optical and Electronic Information, Huazhong University of Science and Technology, Wuhan, 430074, China; Britton Chance Center for Biomedical Photonics, Wuhan National Laboratory for Optoelectronics, Huazhong University of Science and Technology, Wuhan, 430074, China; Department of Bioengineering, University of California, Los Angeles, Los Angeles, 90095, U.S.A.; School of Medicine, University of California, Los Angeles, Los Angeles, 90095, U.S.A.; Joint Department of Bioengineering of UT Arlington/UT Southwestern, University of Texas at Arlington, Arlington 76010, U.S.A.; Chemical and Biomolecular Engineering Department, University of California, Los Angeles, Los Angeles, 90095, U.S.A.; Nanoscience and Nanotechnology Research Center, Research Organization for the 21st Century, Osaka Prefecture University, Osaka, 599-8570, Japan; Mechanical and Aerospace Engineering Department, University of California, Los Angeles, Los Angeles, 90095, U.S.A.

## Abstract

A key challenge when imaging whole biomedical specimens is how to quickly obtain massive cellular information over a large field of view (FOV). Here, we report a sub-voxel light-sheet microscopy (SLSM) method enabling high-throughput volumetric imaging of mesoscale specimens at cellular-resolution. A non-axial, continuous scanning strategy is used to rapidly acquire a stack of large-FOV images with three-dimensional (3-D) nanoscale shifts encoded. Then by adopting a sub-voxel-resolving procedure, the SLSM method models these low-resolution, cross-correlated images in the spatial domain and iteratively recovers a 3-D image with improved resolution throughout the sample. This technique can surpass the optical limit of a conventional light-sheet microscope by more than three times, with high acquisition speeds of gigavoxels per minute. As demonstrated by quick reconstruction (minutes to hours) of various samples, e.g., 3-D cultured cells, an intact mouse heart, mouse brain, and live zebrafish embryo, the SLSM method presents a high-throughput way to circumvent the tradeoff between *intoto* mapping of large-scale tissue (>100 mm^3^) and isotropic imaging of single-cell (~1-μm resolution). It also eliminates the need of complicated mechanical stitching or precisely modulated illumination, using a simple light-sheet setup and fast graphics-processing-unit (GPU)-based computation to achieve high-throughput, high-resolution 3-D microscopy, which could be tailored for a wide range of biomedical applications in pathology, histology, neuroscience, etc.

---

In optical microscopy, high-resolution (HR) volumetric imaging of thick biological specimens is highly desirable for many biomedical applications such as development biology, tissue pathology, digital histology, and neuroscience. To obtain information on cellular events from the larger organism, e.g., a live embryo, intact tissue, or an organ, spatiotemporal patterns from the micro- to meso-scale must be *intoto* determined and analyzed^1-5^. Thus, there is a growing need to develop high-resolution, high-throughput imaging methods that can map entire large-volume specimens at high spatiotemporal resolution^6,7^. Light-sheet microscopy (LSM) has recently emerged as a technique of choice that can image samples with low phototoxicity and at high speed^8-23^. However, similar to conventional epifluorescence methods, LSM remains subject to the fundamental tradeoff between high illumination/detection numerical apertures (NAs) and wide imaging fields of view (FOVs). In addition, accurate digital sampling by the camera is also compromised by the need for large pixel size with high fluorescence sensitivity. Therefore, the achievable resolution of current LSM systems is often pixel-limited under large FOVs, yielding inadequate optical throughput for digital imaging of mesoscale organisms at cellular resolution. Tile imaging-based LSM systems have been developed to artificially increase the space-bandwidth product (SBP)^24^, hence, realizing high resolution imaging of large specimens^18,25-29^. Despite the compromised speed induced by repetitive mechanical stitching, the high illumination/detection NA configuration in tile imaging induces increased phototoxicity for increasing sample size and limits fluorescence extraction from deep tissue. In addition, several techniques such as Fourier ptychographic microscopy^30,31^, synthetic aperture microscopy^32-35^, contact-imaging microscopy^36,37^, wavelength scanning microscopy^38^, and lens-free digital holography^39-41^ have recently provided a computational means of reconstructing a wide-FOV, HR image based on a number of low-resolution (LR) frames having certain correlations in the space, frequency, or spectrum domain^42-44^. However, the majority of these methods target two-dimensional bright-field microscopy and are not compatible with volumetric fluorescence imaging of thick samples.

Here, we present a new imaging method capable of providing three-dimensional (3-D) super-resolution and increased SBP for conventional light-sheet microscopy, without the involvement of mechanical stitching or complicated illumination modulation. This method, termed ‘sub-voxel light-sheet microscopy (SLSM)’, shares its roots with pixel super-resolution (PSR) techniques^36,37,41-43,45^ and works by efficiently computing a number of LR, under-sampled, and shift-modulated LSM volumes in the spatial domain to reconstruct an output with significantly higher resolution throughout the entire sample. Unlike either PSR or conventional LSM, our SLSM method super-resolves large-scale fluorescence images in terms of voxels for the first time, expanding the system SBP, which was originally limited by the low-magnification/NA optics. Furthermore, in SLSM, these special image sequences carrying high-frequency spatial information can be obtained through a simple retrofit of the *z*-scan apparatus employed in conventional LSM systems. Moreover, segmentation of the raw image sequence, modelling of the spatial shifts beyond the system resolution, and reliable estimation of the HR output are all based on a GPU-accelerated parallel computation flow. Therefore, rapid and convenient generation of raw image data, in conjunction with efficient image modelling and reconstruction, realizes high-throughput, HR imaging of large biological specimens via the SLSM technique.

In this study, this SLSM capability is broadly verified by imaging various samples, such as 3-D cultured cells, a live zebrafish embryo, and intact mouse organs. In addition, we demonstrate that SLSM can be combined with multi-view data fusion^46^ to achieve complete imaging of scattering samples, realizing an isotropic resolution of ~1.6 μm (compared to the original ~6.5 and 26 μm) throughout a volume of more than 100 mm^3^. In the following, we elucidate the SLSM experimental setup, discuss implementation of the sub-voxel-resolving (SVR) computation, and demonstrate applications to whole-organism imaging.

## Results

### Off-detection-axis scanning setup

In this study, SLSM imaging was implemented based on a home-built selective plane illumination microscope (SPIM), which is a well-known LSM modality (see Methods and Supplementary Fig. S1 for the full optical layout). This approach functions under a low-magnification setup with a wide FOV covering the entire specimen. Unlike a regular SPIM with *z*-scan parts, the SLSM additionally contains a customized tilting plate that can adjust the scanning axis, allowing sample scanning in a direction with a certain deviation angle *θ* relative to the *z*-axis (detection axis) (Fig. 1a). As the sample is continuously moved across the laser sheet in this non-detection-axis direction, a camera records images of the sequentially illuminated planes at high acquisition speed (Fig. 1b). By matching the frame rate *r* and scanning velocity *v*, a fine step size *s*, typically hundreds of nanometres (*s* = *v/r*), is generated between adjacent frames (Fig. 1b). Thus, every non-axial incremental step provides unit sub-voxel shift components *s_x_, s_y_*, and *s_z_* simultaneously in both the lateral and axial directions, through a one-directional scan (Fig. 1b). In addition, this continuous scanning mode matched by fast camera acquisition provides SLSM with a high imaging throughput of up to hundreds of megavoxel per second. Depending on the sample size and fluorescing intensity, SLSM acquisition usually requires only a few seconds to a few minutes, finally generating a raw image stack containing thousands of spatially modulated frames.

**Figure 1.**
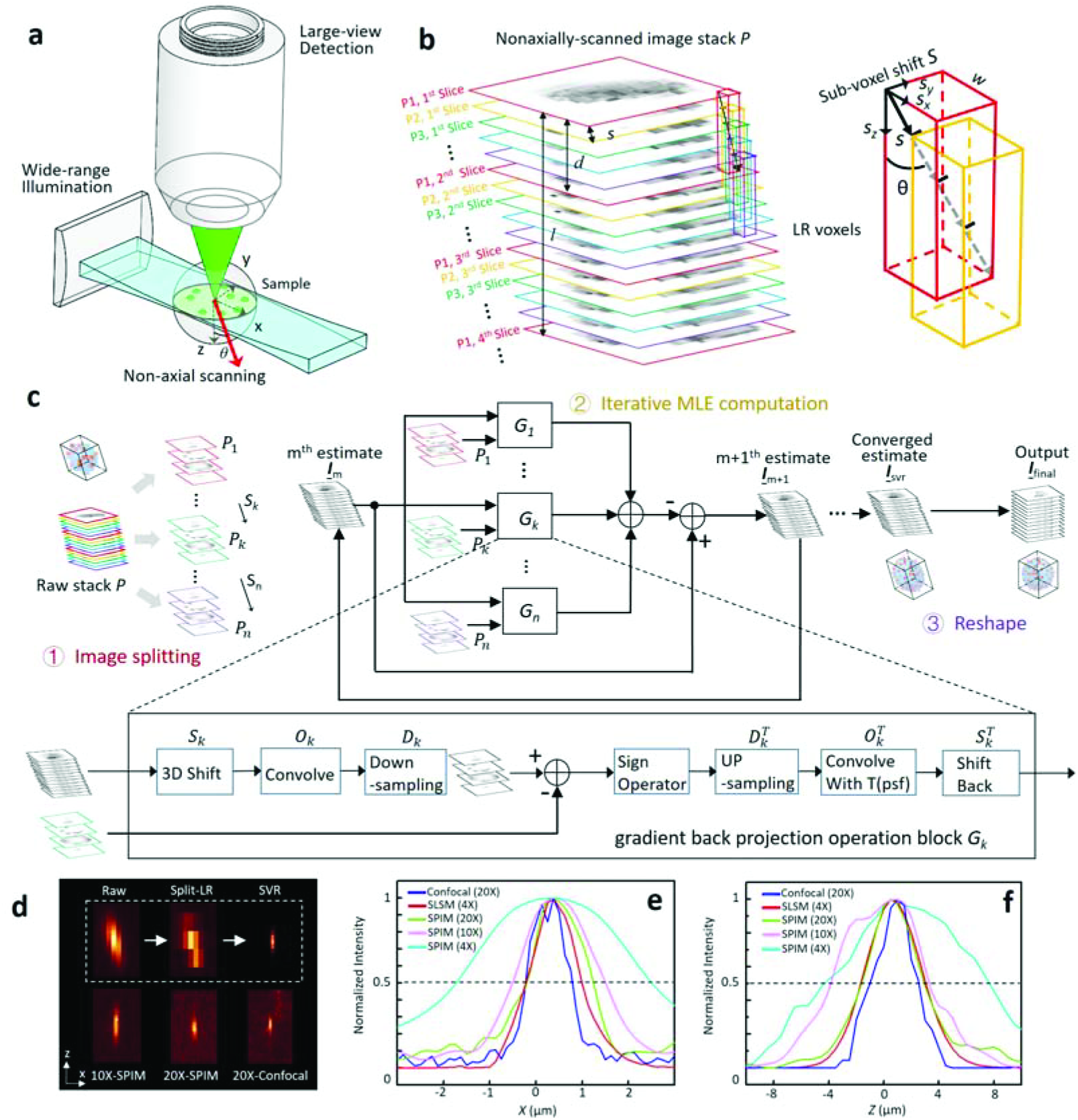
Principles of SLSM. **a**, SLSM geometry with wide laser-sheet illumination and large-FOV detection. The sample is scanned in one direction (red arrow) with the deviation angle *θ* being 10-20° relative to the *z*-axis. **b**, Sequence of spatially modulated, LR images recorded under high frame rate. The step size *s* is much smaller than the laser-sheet longitudinal extent *l* (1/e^2^). The raw image stack is split into multiple sub-stacks (indicated by different colours), with a voxel depth *d* that is one third the laser-sheet thickness *l*. Each LR sub-stack *P_k_* is correlated to the 1^st^ reference stack *P_1_* (red), with a non-axial, sub-voxel shift *s_k_* = (*k* – 1)*\*s*. For instance, *P_2_* (yellow) can be registered to *P_1_* with unit displacement *s*, which consists of three-directional components *s*_x_, *s*_y_, and *s*_z_ simultaneously. **c**, Block diagram of iterative SVR procedure. The spatially correlated, wide-view, LR stacks (step 1) are input to a maximum likelihood estimation (step 2) to iteratively reconstruct a voxel-super-resolved image. Block *G_k_* represents a gradient back-projection operator that compares the *k*th LR image to the estimate of the HR image in the *m*th steepest descent iteration. At the final step (3), voxel re-alignment is applied to recover the sample shape accurately from the slight deformation caused by the tilted scan. **d**, Imaging of 500-nm fluorescent beads using SLSM (4×/0.13-DO/0.022-NA sheet illumination), SPIM (10×/0.3-DO/0.04-NA sheet illumination, 20×/0.45-DO/0.06-NA sheet illumination), and confocal microscope (Zeiss LSM 510, 20×/0.7 objective). The *x-z* planes of the bead are shown to compare the lateral and axial resolving powers of the different methods. **e** and **f**, Intensity plots of linecuts (as shown in **d**) through lateral and axial extents of bead for each method, with 50% intensity level (dashed line) shown for estimation of the FWHM.

### Principles of SVR procedure

An SVR algorithm was designed to specifically segment the acquired shift-encoded, wide-view, LR image stack (denoted by 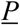) into multiple sub-stacks, model them as probability distributions with subtle spatial correlations, and reconstruct a final output image that encompasses significantly improved resolution over the entire large-volume sample (Fig. 1c). In this approach, the raw image stack 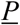 is first divided into a number of LR 3-D images 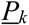 (*k* = 0, 1, 2,…, *n*). In each LR *P_k_*, the voxel width *w* (*x, y*) is simply given by the ratio of the camera pitch size and magnification factor. The voxel depth *d*, denoting the axial spacing implemented when extracting the slices from the raw stack, is set to approximately one third of the laser-sheet longitudinal extent *l* (*1/e^2^*) to satisfy the Nyquist sampling principle. The segmentation is, therefore, performed by re-slicing the raw image every *l/3s_z_* frames, as shown in Figs. 1b and 1c (step 1). The number of total LR images *n*, which implies the amount of sub-voxel information extractable from the raw image, is determined by dividing one LR *w* by *s_x_*; this indicates that these *n* images are correlated with each other in terms of the SVR displacements (Figs. 1b, 1c). Each segmented LR 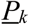 can be considered as a blurred and under-sampled ‘regular SPIM’ image presenting standard resolutions determined by the system optics. Meanwhile, all the *P_k_* can be spatially registered to the HR image 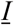 to be solved (also the reference image *P_1_*), according to a non-axial, SVR shift *s_k_* = (*k* – 1)*\*s* (Fig. 1c; red-, green-, and purple-bordered images). The SVR procedure then reconstructs the HR 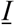 based on computation of these LR images modelled with their known sub-voxel shifts, the system optical blurring, and the camera discretization.

The design of SVR procedure follows a straightforward principle similar to PSR techniques functioning in the 2-D spatial domain^36,42^. It seeks the super-resolved estimate 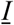 that is most consistent with multiple measurements 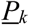 after a series of degradation operators that reasonably models the digital imaging process being successively applied^36,43^. Here, 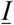 can be specifically solved by minimizing the following cost function:

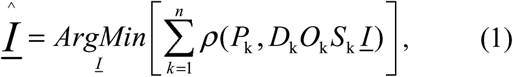

where *ρ* measures the difference between the model and measurements, *S_k_* is the geometric motion operator between the HR estimate 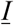 and the *k^th^* LR 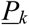, the PSF of the SLSM system is modelled by the blur operator *O*, and *D_k_* represents the decimation operator that models the camera digital sampling. Theoretically, the computation estimates a HR image having maximum likelihood with the LR inputs after given degradations *S_k_, O_k_*, and *D_k_* are applied. Fig. 1c (step 2) shows an iterative maximum likelihood estimation (MLE) solution procedure for 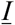. The program first generates an initial guess of HR 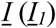, which is simply the interpolation of 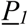. Each *P_k_* is then compared to the warped, blurred, and decimated 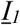 using the gradient back-projection operator (block *G_k_*). Their differences are summed and thereafter weighted by a factor *β*, for calculation of the second estimate. This process is iterated until a converged solution of the *m*th 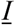 is approached after feeding of the (*m* – 1)th estimate. A voxel re-alignment is applied in the final step to recover an accurate reconstruction from the slight deformation caused by the non-axial scan (Fig. 1c, step 3). Detailed implementation of the SVR procedure is elucidated in the Supplementary Notes and Supplementary Fig. S4, and simulation results are shown in Supplementary Fig. S6. Optimization of the SVR parameters is discussed in the Supplementary Notes and Supplementary Figs. S5.

### Imaging characterization

Simulation results of the proposed SVR procedure are provided in Supplementary Fig. S6, whereas Figs. 1d-1f experimentally characterize the SLSM performance through imaging of fluorescent microbeads (~500-nm diameter) under a 4×/0.13 detection objective (DO) plus 0.022-NA plane illumination (PI) configuration. The microbeads were scanned along a *θ* of 10° (to the *y-z* and *x-z* planes) with an *s* of 144 nm, yielding an incremental lateral shift *s_x_* and axial shift *s*_y_ of 25 and 140 nm, respectively. Then, 65 groups (*n*) of LR, 3-D images (voxel size: 1.625 × 1.625 × 6 μm^3^) were extracted from the raw sequence to compute an SVR image with fourfold enhancement in each dimension. The LR and SVR results were compared to mid-magnification SPIM (10×/0.3-DO/0.04-NA PI), high-magnification SPIM (20×/0.45-DO/0.06-NA PI), and confocal microscope (20×/0.7 objective) images, as shown in Fig. 1d. The line intensity profiles of the resolved beads are plotted in Figs. 1e and 1f, to compare the lateral and axial resolutions of these methods. The achievable lateral and axial full widths at half maximum (FWHMs) of the 4×-SLSM were improved from ~4 and 13 μm to ~1.2 and 4.5 μm, respectively, being similar to the results for the 20×-SPIM and 20×-confocal microscope.

We validated the SLSM by obtaining HR reconstructions of 3-D-cultured normal human bronchial epithelial (NHBE) cell spheroids (labelled with DAPI nucleic acid staining, see Methods section), as shown in Fig. 2a. A sequence of images from consecutively illuminated planes was rapidly recorded in 1 min under a 100-fps acquisition rate, targeting a specific volume-of-interest containing dense cell spheroids (Fig. 2a). Note that each LR image subdivided from the raw stack simply adopts the limited resolution from the system optics; hence, only the rough shape and distribution of the selected spheroid were identified, whereas the single cells remained unresolvable (Fig. 2b1). The SVR procedure then began with an initial guess, which was simply an ×4 interpolation of the first reference image (Fig. 2b2), and iteratively converged to the final HR image, where the chromatins inside the cell nucleus became distinguishable (Fig. 2b3). Indeed, the linecuts through the cell spheroids (Fig. 2d) by each method revealed substantially improved resolution from SVR, which surpassed 20 × -SPIM result with observable spherical aberration shown (Fig. 2c, Supplementary Fig. S3).

**Figure 2.**
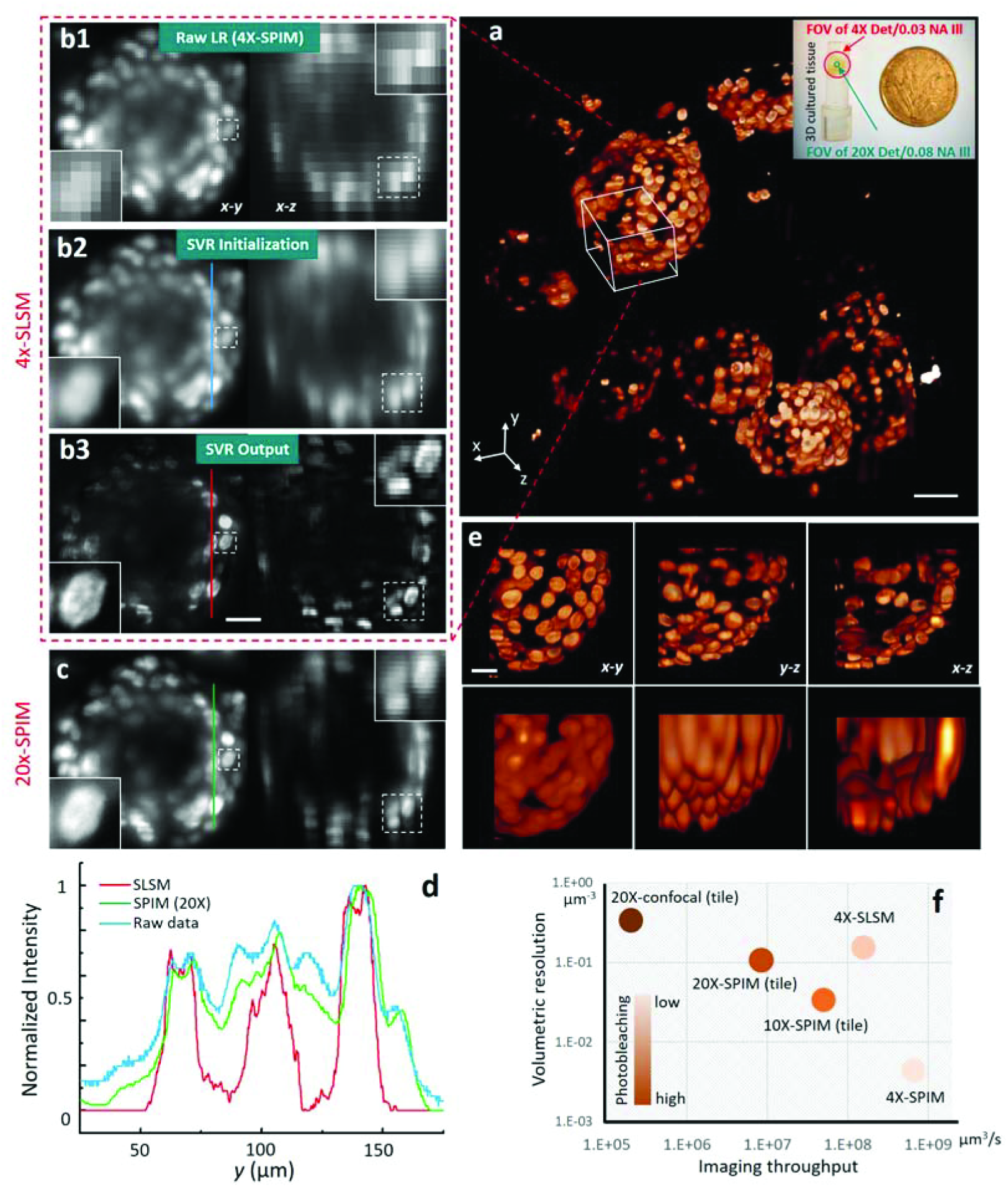
SLSM demonstration of 3-D-cultured NHBE cells. **a**, NHBE cell spheroids three-dimensionally cultured in a piece of Matrigel substrate. The cells were fluorescently labelled through DAPI nuclear staining for SLSM reconstruction. **b1**, LR raw image of selected cell spheroid (first of 64 groups), captured under illumination and detection NAs of 0.022 and 0.13, respectively, with 1.625 × 1.625 × 6 μm^3^ voxel size. The vignette views show a single-cell nucleus. **b2**, Initial estimate of SVR computation, which is an ×4 interpolation of b1. **b3**, Final SVR reconstruction of the same region, with a reconstructed voxel size of 0.41 × 0.41 × 1.5 μm^3^. The features of the cell nucleus were recovered through blurring and pixelation, with the chromatin becoming discernible. **c**, Images of same cell spheroid taken by SPIM using 20×/0.45 objective and 0.06-NA illumination, for comparison. **d**, Intensity plot of linecuts (shown in **b** and **c)** for each setting, indicating that SVR provides substantively improved contrast and resolution. **e**, Volume renderings in raw mode and SVR reconstruction, comparing 3-D localization accuracy of single cells in self-assembled spheroid. Scale bars: 20 μm (5 μm in insets). **f**, Comparison of imaging resolution, speed, and photobleaching for 4×-SPIM, 10×-SPIM tile imaging, 20×-SPIM tile imaging, 20x confocal microscopy, and 4×SLSM. The achieved effective throughputs are 2.7, 1.7, 0.9, 0.07, and 24 megavoxel SBP per second, respectively.

Besides the selective image planes, we compared the volume rendering of a reconstructed cell spheroid given by conventional 4×-SPIM and 4×-SLSM, with the latter explicitly showing much improved 3-D single-cell localization (Fig. 2e). SLSM provides a wide FOV of ~23 mm^2^ from its low-magnification DO (4×/0.13) and low NA PI (0.022), whereas its achieved resolutions are similar to those of 20×-SPIM (20×/0.45-DO/0.07-NA PI). Thus, the SLSM can be regarded as a light-sheet microscope combining the FOV advantage of 4x-SPIM with the resolution advantage of 20×-SPIM. From another perspective, its stitching-free, continuous scanning mode exhibits a much higher acquisition throughput as well as lower photo-bleaching than the stitching results of the 10×-SPIM, 20×-SPIM, and confocal microscope (Supplementary Notes).

We calculated the imaging resolution, speed, and photobleaching of 4×-SPIM, 10×-SPIM tile imaging, 20×-SPIM tile imaging, 20×confocal microscopy, and 4×-SLSM. As indicated in Fig. 2f, the SLSM exhibited the highest effective throughput at ~25 megavoxel SBP per second, which is more than 10 times higher than those of the other modalities. Besides the increased SBP (FOV divided by resolution), the SVR computation, to some weak fluorescing extent, improves the signal-to-noise ratio after multiple image fusion (Supplementary Fig. S7). Furthermore, circumventing the use of high-NA, high-maintenance optics renders SLSM considerably less vulnerable to chromatic aberration occurring in multi-colour illumination (Supplementary Notes and Supplementary Fig. S2), and resistant to the spherical aberration that causes severe image deterioration for deep tissue (Supplementary Fig. S3). In the applications described below, this underlying robustness allows the SLSM prototype to image thick specimens at high spatial-temporal performance while retaining a relatively simple setup.

### Multi-colour 3-D whole-organism imaging at high-throughput

SLSM does not require special engineering of the fluorescence emission to generate image modulation, rendering it compatible with various labelling techniques. In Fig. 3, we demonstrate SLSM imaging of two types of specimen: an optically cleared intact mouse heart (neonate, D2) exhibiting endogenous autofluorescence in cardiomyocytes (Fig. 3a), and a two-colour transgenic zebrafish embryo (3 days post fertilization, d.p.f.) tagged with green and red fluorescence protein at the motor neurons (Islet1-GFP) and somite fast muscles (mlcr-DsRed), respectively (Fig. 3b). Furthermore, a time-course study on an anaesthetized live fish embryo from 48 to 72 hours post fertilization (h.p.f.) was implemented in this work to investigate the neuron/muscle development in the hinderbrain region, as shown in Supplementary Fig. S9. The raw data acquisition was very fast, at a rate above 200 megavoxels per second.

**Figure 3.**
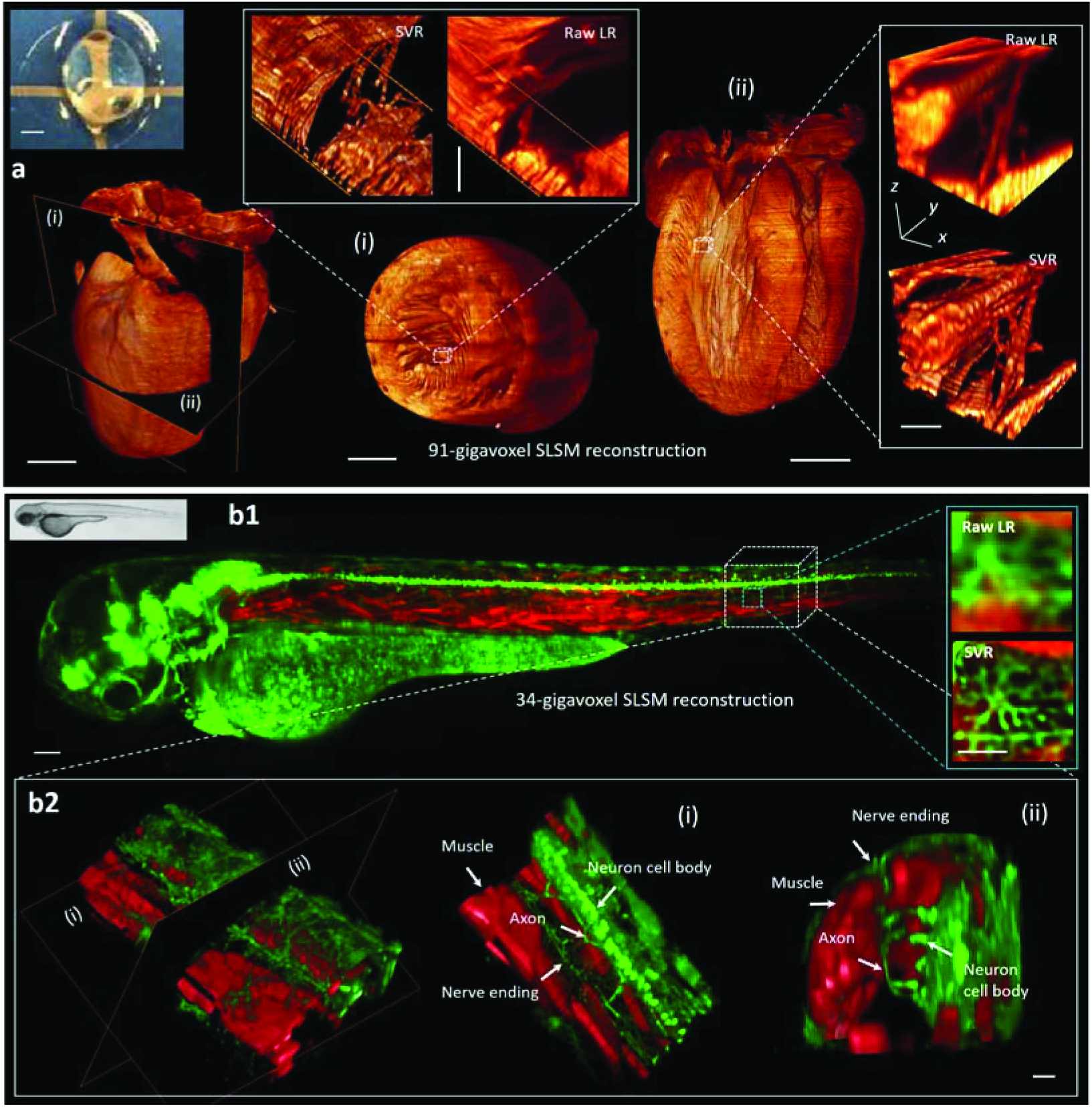
Rapid high-throughput volumetric imaging of whole organisms via SLSM. **a**, 91-gigavoxel SVR reconstruction of optically cleared intact mouse heart (inset: neonate day 1) with endogenous autofluorescence. The magnified views of the slices along the indicated coronal and axial planes (i and ii) reveal that the SVR procedure substantively improves visualization of the cardiomyocytes compared to the raw images. The image acquisition and SVR reconstruction time for the entire heart were approximately 4 and 12 min, respectively. Scale bars: 1 mm (magnified views: 50 μm). **b**, Two-colour, 34-gigavoxel SVR reconstruction of anaesthetized live zebrafish embryo (3 d.p.f.) tagged with Islet1-GFP at motor neurons and mlcr-DsRed at somite fast muscle. **b2**, Magnified volume rendering in SVR reconstruction of fish somite section, elucidating the interplay of the motor neurons with the muscles during development. The right-side images show slices through the somite along the planes shown in the image on the left. The SLSM image acquisition was completed in 120 s for dual colours, followed by 200 s of GPU-based processing. Scale bars: 100 μm for the whole embryo image (magnified views: 20 μm).

Meanwhile, to match the high-speed acquisition, we developed a GPU-based parallel computation flow to greatly accelerate the SVR procedure simultaneously, super-resolving the large-scale data at a high processing throughput above 100 megavoxels per second. Hence, SLSM can rapidly image (acquisition + computation) these millimetre-size organisms on a scale of dozens of gigavoxel, providing super-resolved cellular structure-function information, such as myocardium architectures and interactions of developing motor neurons with somite muscles, within a few minutes to an hour. In Supplementary Figs. S10 and S11, broader demonstrations of SLSM applications are provided, for whole organism imaging of a Tg (cmlc2:GFP) adult zebrafish heart (60 d.p.f.) and an *αMHC^Cre^; R26^VT2/GK^* murine heart (three colours, P1), which are spatially or temporally more challenging to regular light microscopes.

### Multi-view SLSM for isotropic super-resolution imaging

Despite the use of chemical clearing, light scattering from deep tissue continues to pose a challenge for optical microscopy. Both laser excitation and fluorescence emission experience deflection and attenuation, which together deteriorate signals severely. In addition, even light-sheet setup remains anisotropic for the imaging of mesoscale samples, showing suboptimal axial resolution for several applications such as cell phenotyping and neuronal tracing^18^. The multi-view fusion method^46,47^ was previously developed to address these problems. It functions by registering, weighting, and fusing a number of image stacks recorded under different views, and finally recovers a stack that shows complete signals with improved axial resolution.

Here, we demonstrate that the SVR procedure can be combined with a multi-view approach to *intoto* image thick and scattering samples at near isotropically improved resolution. We developed a 4-view SLSM prototype by imaging a human umbilical vein endothelial cell (HUVEC) and human dermal fibroblast (HDF) 3-D sprouting network (Methods). The cells were co-cultured in a fibrinogen-based hydrogel mixed with dextran-coated Cytodex 3 microbeads (SiO2), finally forming a complex and light-scattering cellular network containing fibrinogen, HUVEC beads, and HDF cells.

First, the sample was rotated by 90° for each set of acquired 2×-SLSM data (2×/0.06 detection, 0.015-NA sheet illumination, 580-nm step size, 7000 frames in 140 s), with four views of the raw stacks being obtained in total. Each set was sub-voxel-resolved separately based on 32 groups of segmented LR inputs, generating four views of the anisotropic SVR image (4 ×4 × 2 enhancement). Multi-view registration followed by weighted fusion was included in the final step to produce an output image with isotropic SVR. This multi-view SVR (mv-SVR) workflow is illustrated in Fig. 4a.

**Figure 4.**
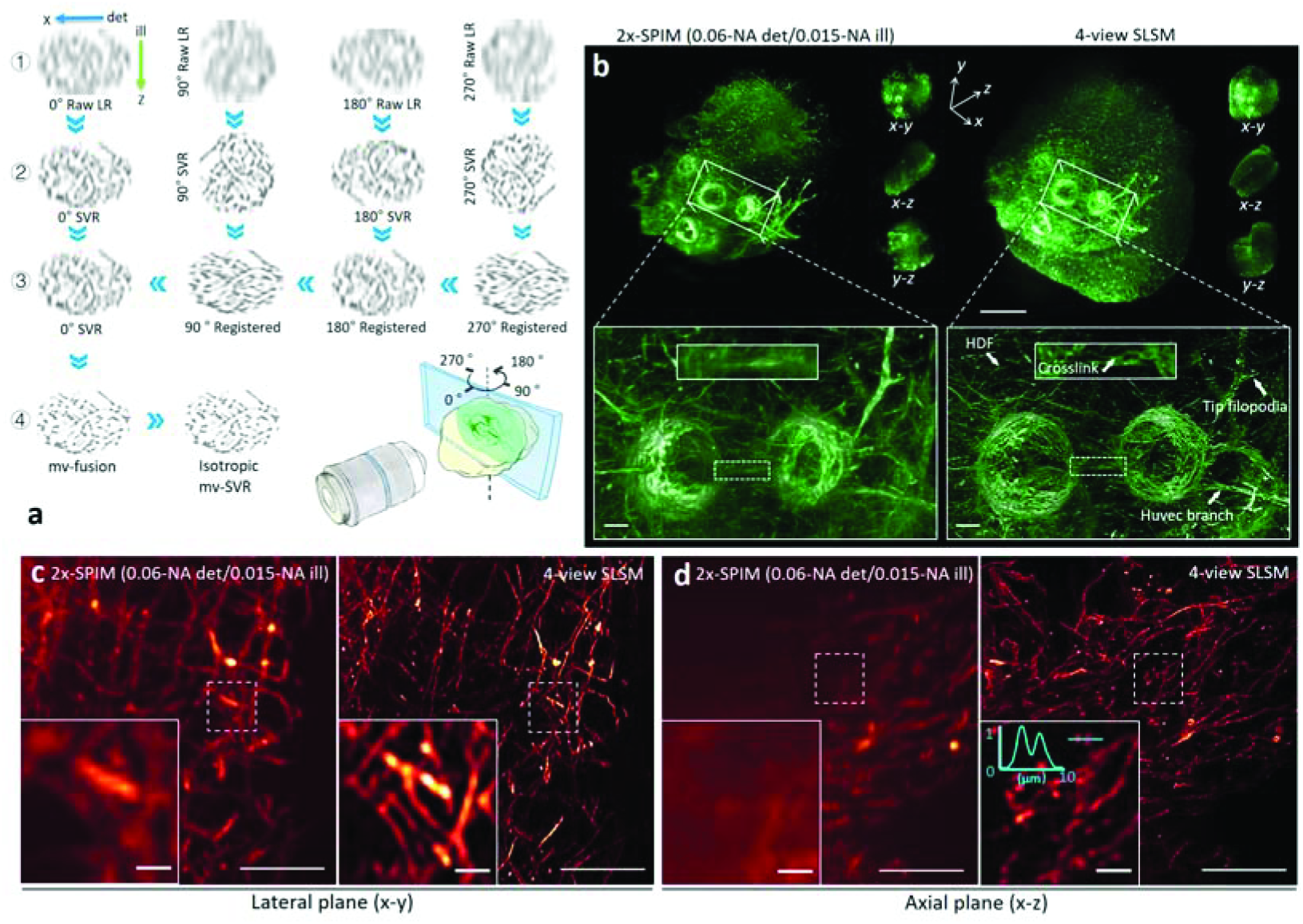
mv-SLSM for three-dimensional isotropic imaging of large scattering HUVEC-HDF cell sprouting network. **a**, The mv-SVR procedure involves the following steps: 1. image acquisition under multiple views; 2. SVR computation for each view; 3. image registration in 3-D space; 4. weighted mv-SVR fusion with final deconvolution. **b**, 3-D reconstruction of cells obtained using single-view SPIM (2×/0.06-DO/0.015-NA sheet-illumination) and 4-view SLSM using the same optics. The SPIM system SBP is 1.1 gigavoxel for the entire sample, whereas 4-view SLSM creates a 190-gigavoxel SBP that renders a higher-resolution and more complete sample structure across a large FOV of ~100 mm^3^. The vignette HR views compare visualizations of the HUVEC branch, tip filopodia, and HDF co-localization (white arrowheads). **c, d**, Maximum intensity projections of supporting HDF cells in *x-y* and *x-z* planes. mv-SVR reconstruction enables clear identification of single fibre sprouting (right columns of **c, d**), which remains very fuzzy in the single-view SPIM image (left columns of **c, d**). The intensity plot of the blue linecut (inset, **d**) indicates a small resolvable distance of ~1.6 μm. Besides isotropic resolution enhancement, mv-SVR reconstruction also restores the highly scattered area that is originally subject to severe signal loss in the single-view image (left area in **d**). Scale bars: 1 mm in **a, b** (magnified views: 100 μm) and 100 μm in **c, d** (magnified vignette views: 10 μm).

In Fig. 4b, we compare the reconstructed volume renderings of a 0° SPIM image (1.1 gigavoxels, voxel size: 3.25 × 3.25 × 9 μm^3^) and 4-view SLSM (190 gigavoxels, reconstructed voxel size: 0.81 × 0.81 × 0.81 μm^3^). Because of the strong scattering of the fibrinogen-microbead substrate, the single-view SPIM visualizes an incomplete structure with poor resolution (Fig. 4b, left). In contrast, the 4-view SLSM reconstructs the entire sample, showing details of HUVEC sprouts from two adjacent dextran beads (Fig. 4b, right). HR local views containing dense HDF fibres are given in Figs. 4c and 4d, also compared to single-view SPIM results. Besides the isotropically improved resolution (resolvable distance: <1.6 μm; Fig. 4d, inset) that enables clear identification of single sprouting HDF cells, the multi-view SLSM (mv-SLSM) also restores the highly light-scattered area in the reference view (Fig. 4d, left). Note that multi-view sub-voxel computation relies on the image registration and weighted fusion in Fourier space and, therefore, the processing speed of 4-view-fused SVR is approximately one order lower than the computation for a single view, reconstructing 18 megavoxels per second on average. Finally, by creating an isotropic, HR, panoramic visualization that encompasses 190 gigavoxels across a volume exceeding 100 mm3 (total processing time: ~3 h), the mv-SVR procedure can accurately analyze vast numbers of cells, such as sprouting branches and tip filopodia, over a piece of mesoscale tissue. Hence, mv-SVR provides a solid foundation for exploration of mature vessel formation and vessel anastomosis processes, both of which are crucial for studying tissue regeneration.

### Fast quantitative mapping of mouse-brain neuronal networks

We reconstructed a P30 Thy1-GFP-M mouse brain using 8-view SLSM (Fig. 5a). HR volumetric renderings of the hippocampus (i), thalamus (ii), and cortex (iii) regions are shown in Figs. 5b-5d, respectively. Compared to conventional SPIM with suboptimal quality, the 8-view SLSM imaging resolved fine neuronal sub-structures such as dendrites and axons at an isotropic resolution of ~1 μm (350-gigavoxel SBP). Benefitting from remarkably improved visualization, neuronal subsets such as clustered astrocyte neurons in the thalamus region could be segmented with clearly separated nerve fibres (Fig. 5e, Imaris software). Three long-projection neurons were also successfully identified and registered in the P30 reference mouse brain. The pathways of these projection neurons were subsequently annotated according to the standard mice brain atlas^48^, as shown in Figs. 5e (ii-iv). The high-resolution, high-throughput features of SLSM enable fast and accurate tracing of both long-distance projections and pathways within the neuronal circuits in the brain of a fluorescent protein transgenic mouse.

**Figure 5.**
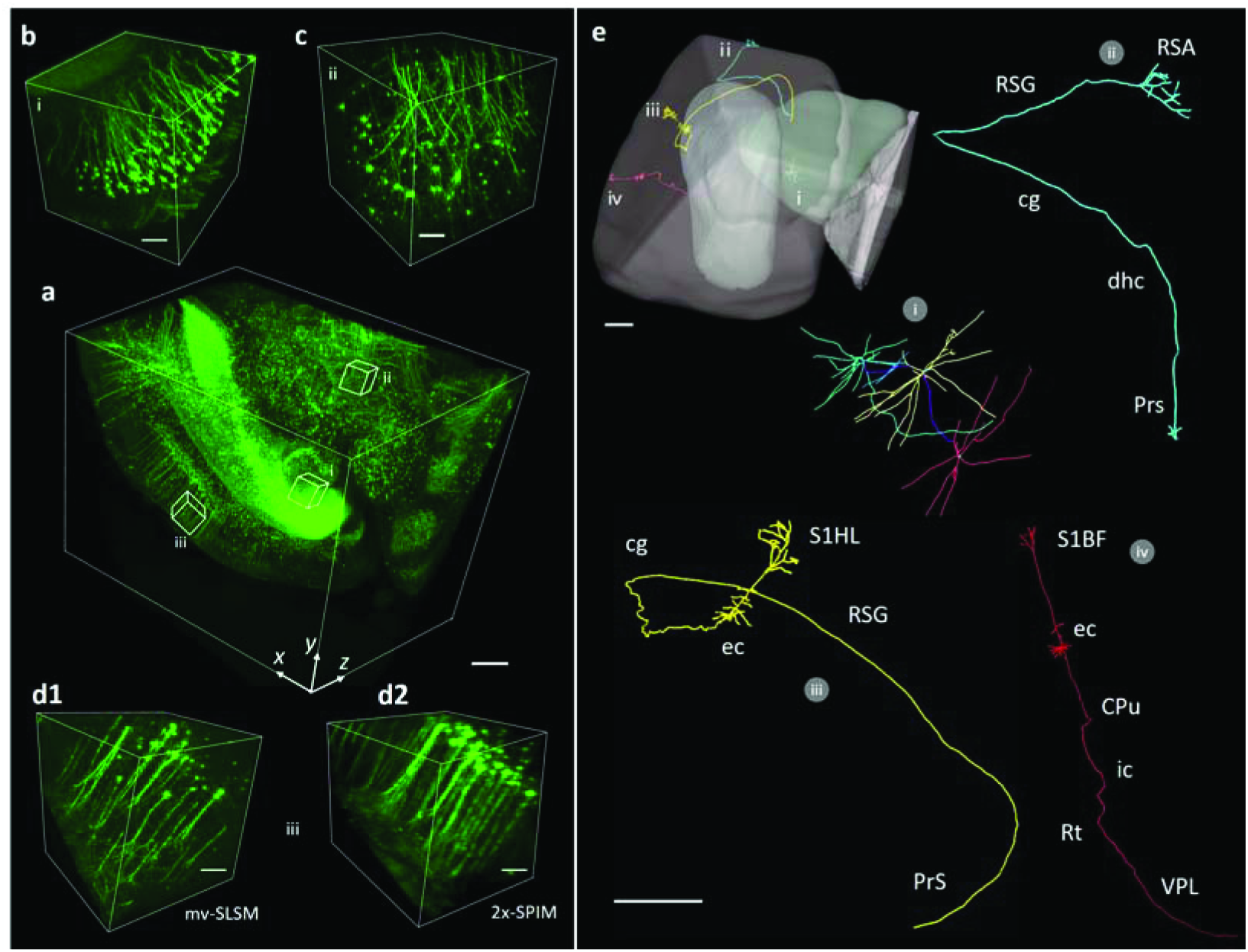
Three-dimensional neuronal mapping of adult mouse brain via 8-view SLSM. **a**, Isotropic SVR volume rendering of optically cleared Thy1-GFP-M mouse brain block (350-gigavoxel scale; scale bar: 500 μm). **b, c, d1**, HR volumetric views of hippocampus, thalamus, and cortex regions, respectively. **d2**, Cortex reconstructed using regular SPIM with 4×/0.13-DO plus 0.022-NA PI, for comparison (scale bar: 100 μm). **e**, Analysis of neural structures and connectivity. Three clustered astrocyte neurons in the thalamus region are segmented with different colours encoded in (i). Three registered projection neurons across the brain block are annotated according to the mice brain atlas (ii, iii, and iv). Abbreviations: cg: cingulum; ec: external capsule; dhc: dorsal hippocampal commissure; CPu: caudate putamen; ic: internal capsule; Rt: reticular thalamic nucleus; Prs: presubiculum; RSG: retrosplenial granular cortex; S1HL: primary somatosensory cortex hindlimb; RSA: retrosplenial agranular cortex. Scale bar: 500 μm.

## Discussion

SLSM is based on unconventional off-axis scanning together with sub-voxel reconstruction, so as to computationally surpass the resolution limit of a regular light-sheet microscope. This technique can be applied to most existing light-sheet microscopes by retrofitting with a readily available tilting stage, thereby expanding the optical throughput for fast high-resolution mapping of large biomedical specimens. In principle, the SVR procedure estimates a 3-D image that best suits a certain conditional probability in the spatial domain. Provided the aperture function and sub-voxel motion are accurately characterized, the maximum-likelihood link between the actual sample profile and recorded data allows the SVR to iteratively render a high-resolution image using a limited-NA, under-sampled configuration, which was originally incapable of providing such a small PSF (Fig. 1).

Unlike most super-resolution fluorescence microscopy methods, which image a single cell beyond the diffraction limit by processing multiple frames acquired under a large NA using highly specialized optics, SLSM is free from either illumination modulation or particular fluorescence labelling, being designed to rapidly enhance the inadequate accuracy obtained by unravelling large organisms under an ordinary small-NA/large-view configuration. Its stitching-free, high-speed image acquisition followed by a parallelized GPU processing flow provides super-resolved 3-D visualization at quasi-real-time throughput. In this work, by imaging a variety of biological specimens from 3-D cells to developing embryos and intact organs, our SLSM robustly exhibited improved performance at low hardware and time cost, using efficient computation to circumvent the trade-off between larger imaging volume and more resolvable detail. The ability to rapidly accomplish cellular imaging of mesoscale organisms at hundreds-of-gigavoxel SBP renders SLSM a valuable tool for wide biomedical applications such as phenotype screening, embryogenesis and brain connectome in histology, development, and neuroscience research, for which both large-scale statistics and high-resolution details are highly desired.

In addition to single-view mode, SLSM can also be expanded by combination with multi-view fusion. Through a rational balance between higher throughput and increased views, mv-SLSM can image thick and scattering samples with isotropic super-resolution and at moderately high throughput. More broadly speaking, this method is currently optimized for light-sheet microscopy, which has a relatively high image contrast and fast acquisition rate. However, the nonaxial scan as well as the SVR computation may be equally suited to other 3-D microscopy methods, such as confocal microscopy and 3-D deconvolution microscopy. Furthermore, we believe this sub-voxel imaging strategy could prove to be transformative, as it provides a general iterative resolution recovery method that could potentially be employed to improve other 3-D imaging modalities, which are possibly limited by inadequate sampling and poor focusing capability.

## Methods

### Experimental setup

A four-wavelength, fibre-coupled semiconductor laser (CNI Laser, RGB 637/532/488/405, China) was used as an excitation source. The laser was first transformed into a collimated Gaussian beam with diameter ~8 mm (1/*e*^2^ value). Then, an adjustable mechanical slit (0-8-mm aperture) was used to truncate the beam in the horizontal direction and thereby tune the aperture of plane illumination. The illuminating cylindrical lens (focal length *f* = 40 mm) finally formed a wide laser sheet that optically sectioned large specimens with an NA adjustable from 0 to 0.1. Then, a ×2 or ×4 (Nikon Plan Apo Fluor 2×/0.06 or 4×/0.13 objective) infinity-corrected, wide-field detection path was constructed orthogonally to the PI path to collect the fluorescent signals. A four-degree-of-freedom motorized stage (*x, y, z* translation and rotation around the *y*-axis, Thorlabs) integrated with a pair of customized tilting plates (10° inclined surface) was constructed for sample mounting and scanning across the laser sheet in an off-detection-axis direction (Supplementary Fig. S1). For a given imaging configuration, the choice of *θ* was balanced between generating a small lateral shift component for a larger enhancement factor and scanning a short axial distance when completing an entire voxel shift. Usually, the diagonal of the LR voxel is chosen as the nonaxial scanning axis, with *θ* being 10-15°. A scientific complementary metal-oxide semiconductor (sCMOS) camera (Hamamatsu Orca Flash 4.0 v2 or Andor Zyla 5.5, pitch size: 6.5 μm) continuously recorded the planar images from the consecutively illuminated planes at a high speed of up to 200 frames per second.

### SPIM, SLSM, and multi-view SLSM acquisition

For the conventional SPIM imaging, the sample was scanned along the *z*-axis with 2–9-μm step size depending on the different illumination NAs. To avoid motion blur generated by these incremental moves of several microns, it was necessary to implement the *z*-scan in a step-by-step manner by synchronizing the laser, motor, and camera using LabVIEW software (version 2014, National Instrument, US.A.). This stepwise *z*-scan limited the acquisition speed to a maximum rate of 13 fps in our system (Thorlabs stepper ZST225B). Further, 10× and 20× tile SPIM imaging were automatically implemented in a *z-x-y* raster scanning method, and the acquired mosaic volumes were subsequently stitched together using the Grid/Collection stitching plugin (ImageJ software). Each time, only a small region of the entire illuminated plane was imaged; thus, the tile imaging was subjected to a photobleaching rate higher than those for the SPIM and SLSM.

For SLSM imaging, the camera was synchronized with a continuous sample scan under LabVIEW control, recording images in sequential mode. The step size could, thus, be determined by matching the scanning velocity (10-30 μm/s) and camera frame rate (20-200 fps). Depending on the sample dimension, tilt angle, and enhancement factor, this value varied from 140 to 600 nm. Compared to the voxel size of several microns in the segmented LR volumes, the motion blur caused by the continuous scan was negligible under such a small step size and did not affect the SVR computation accuracy. At the same time, the SLSM image acquisition could be fast under continuous mode, taking 30 to 300 s to obtain a sequence containing thousands to tens of thousands of frames. For multi-view imaging, the stepper rotated the thick and scattering samples four to eight times, acquiring a number of raw SLSM image stacks under different views. The data stream was transferred from the camera to a RAID 0 volume of solid state drives (4 × Crucial M550 1TB) in real time. The raw images were finally saved in 16-bit TIFF format on the gigavoxel to tens of gigavoxel scale.

### Efficient GPU-accelerated SVR reconstruction

In practice, a steepest descent method was used in the SVR computation, to iteratively approach a converged super-resolved solution at high efficiency. Here,

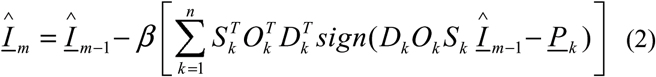

where 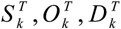 represent the inverse operations of *S_k_,O_k_, D_k_*, respectively. A Matlab script was first developed as a proof of principle for the SVR (Supplementary Notes and Supplementary Fig. S4). Using a desktop workstation with dual Intel E5-2630 v4 central processing units (CPUs) (no GPU involved), the processing throughput did not exceed 0.3 megavoxels per second, almost three orders slower than the image acquisition.

Reconstructing data with volumes of tens to hundreds of gigavoxels for large specimens would require an impractically long time, i.e., multiple days. Therefore, we developed a GPU-based parallel workflow to accelerate the processing, achieving near three-order acceleration (Supplementary Table 1). Depending on the degree of parallelization and the number of views provided, the processing throughput varied from tens to hundreds of megavoxels per second, depending on the power of the dual NVidia TESLA P100 graphical cards. For example, in single-view configuration, the program explicitly finished a HR reconstruction of an intact heart in ~12 min (90 gigavoxels), and a dual-colour reconstruction of an entire zebrafish embryo in ~4 min (34 gigavoxels). It should be noted that this speed could be further increased by employing additional GPUs.

For multi-view SVR of thick and scattering HUVEC–HDF sprouting, 32 groups of LR images were extracted from the raw sequence of each view. The unit lateral and axial shifts were 100 and 558 nm, respectively. Considering the optical blurring and camera under-sampling, the effective lateral and axial resolutions of each LR image were approximately 7 and 26 μm, respectively, yielding ~1.1-gigavoxel SBP (voxel size: 3.25 × 3.25 × 9 μm^3^) over ~100-mm^3^ volume.

Applying the aforementioned SVR procedure, the super-resolved image *I_single_* obtained under each view could be reconstructed as intermediate results. In each reconstructed *I_single_*, an increased SBP of 34 gigavoxels was presented anisotropically (0.81 × 0.81 × 4.5 μm^3^ voxel spacing). According to the reported multi-view fusion method, we interpolated all *I_single_* with isotropic 0.81-μm pitch size, rotated the second view and registered it to the 0° reference view by iteratively matching their histograms. An initial fusion of the images was generated by taking the weighted average of the two registered views in Fourier space:

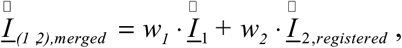

where 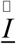 represents the Fourier transform of 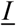 and the *w* terms are the corresponding weights that reasonably average the 2 SVR views. Then, this initial fusion was used for the second registration and fusion iteration, in which the reference became the fusion of the first stage. By repeating this process, we obtained the fused image *I_merged_* that accumulated the effective information from all the SVR views^46^. This procedure can be expressed as follows:

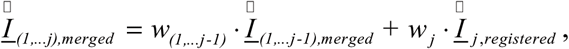

where *j* = 3,4 in a four-view SVR fusion. The weighting of the average was determined by the expected signal-to-noise ratio as follows:

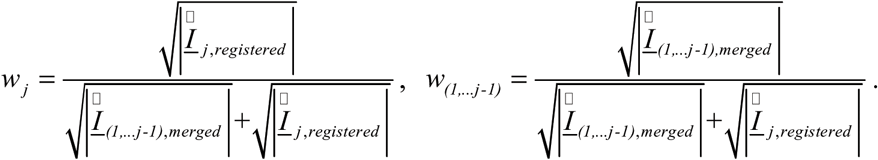

A multi-view deconvolution was applied in the last step to obtain the deblurred output, which exhibited a final SBP of ~190 gigavoxels with isotropically improved resolution. Implementation of the SVR and mv-SVR methods is also detailed in the Supplementary Notes. At present, as a proof-of-concept, the mv-SVR is based on histogram registration and weighted fusion, and has not been performance optimized. Note that more efficient bead-based registration^47^ and Bayesian deconvolution^49^ have recently been reported. We believe SVR can be combined with these techniques to yield even better image quality and higher speed. Further, the capacity was verified using four-view data in this work; however, it is quite certain that the result can be further optimized through provision of additional views^46^.

### Image visualization and analysis

The visualizations of SPIM, SLSM, and confocal data were performed using Amira (Visage Imaging). Planar images were presented in their original formats, unless otherwise mentioned, with no sharpening, interpolation, or registration applied. Maximum intensity projections and volume renderings were performed using the ProjectionView and Voltex functions in Amira with built-in colourmaps. Long-projection neuronal tracing of the clarified mouse brain was performed using the Filament module in Imaris (Bitplane). See the Supplementary Figures for additional details and spatiotemporal visualizations of the developing zebrafish embryo, adult zebrafish fish heart, and neonate mouse heart.

### Preparation of large-scale, 3-D-cultured cells

NHBE cells (Lonza, Walkersville, MD) were prepared. The growth factor reduced reconstituted basement membrane known as Matrigel (BD Biosciences, Bedford, MA) was used for the 3-D culture experiments. The cells were seeded into 300 μl of Matrigel in liquid state and gently mixed with a pipet. The cells were then placed in a 24-well plate and incubated for 25 min at 37 °C to gelatinize. Next, 1 ml of bronchial epithelial growth medium (BEGM; Lonza, Walkersville, MD) was added to the 24-well plate and the medium was exchanged every other day. **Fluorescent staining.** Direct and indirect immunofluorescent staining was applied to the cells to visualize the branching structure. For fixation, 4 % paraformaldehyde (Electron Microscopy Science, Hatfield, PA) was applied to the Matrigel at room temperature for 20 min. After washing with phosphate buffered saline (PBS), PBS containing 0.5% Triton X-100 was applied for cell permeabilization for 10 min at 4 °C. This was followed by three 10-min washes with PBS. Then, the gels were blocked with 10% goat serum and 1% goat anti-mouse immunoglobulin G (Sigma-Aldrich, St. Louis, MO) in IF buffer (0.2% Triton X-100; 0.1% bovine serum albumin (BSA) and 0.05% Tween-20 in PBS). As a primary antibody, rabbit E-cadherin monoclonal antibody (Life Technologies, Grand Island, NY) was incubated with Matrigel overnight at 4 °C and the gel was rinsed three times with 10% goat serum for 20 min each. Then, Alexa Fluor 555 goat anti-rabbit IgG was incubated for 2 h at room temperature followed by three 20-min rinses with PBS. For nuclear staining, DAPI was incubated for 20 min at room temperature followed by three 20-min rinses.

HUVECs were mixed with dextran-coated Cytodex 3 microcarriers at a concentration of 400 HUVEC per bead in 1 ml of endothelial cell growth medium (EGM)-2. The beads with cells were shaken gently every 20 min for 4 h at 37 °C and 5% CO_2_, and then transferred to a 25-cm^2^ tissue culture flask and left for 12–16 h in 5 ml of EGM-2 at 37 °C and 5% CO_2_. The following day, the beads with cells were washed three times with 1 ml of EGM-2 and re-suspended at a concentration of 500 beads/ml in 2-mg/ml fibrinogen, 1-U/ml factor XIII, 0.04-U/ml aprotinin, and 80,000-cell/ml HDF at a pH of 7.4. Then, 250 μl of this fibrinogen/bead solution was added to 0.16 units of thrombin in one well of each of the glass-bottom 24-well plates. The fibrinogen/HUVEC bead/HDF cell solution was allowed to clot for 5 min at room temperature and then at 37 °C and 5% CO_2_ for 20 min. EGM-2 was added to each well and equilibrated with the fibrin clot for 30 min at 37 °C and 5% CO_2_. The medium was removed from the well and replaced with 1 ml of fresh EGM-2 and was later changed every other day. The co-culture assays were monitored for 7 days and then fixed and stained with Alexa Fluor 488 for imaging of the cytoskeletons.

### Zebrafish embryo culture

Transgenic zebrafishes (Islet1:GFP-mlcr:DsRed and cmlc2:GFP) were raised in the zebrafish core facility of the University of California, Los Angeles (UCLA). All experiments were performed in compliance with the approval of the GLA Institutional Animal Care and UCLA Institutional Animal Care and Use Committee (IACUC) protocols. To maintain transparency of the zebrafish embryos, they were incubated with egg water containing 0.2-mM 1-phenyl-2-thio-urea (Sigma) to suppress pigmentation at 24 h.p.f. The live fish embryos were anaesthetized with low-concentration tricaine (0.04 mg/ml, MS-222, Sigma) before being mounted in a fluorinated ethylene propylene (FEP) tube for sustained imaging.

### Optical clearing of thick organs

The adult mouse brain (Thy1-GFP-M), neonate mouse heart (wild-type, αMHCCre; R26VT2/GK), and adult zebrafish heart (cmlc2-GFP) were originally turbid organs. As a result, it was necessary to perform tissue optical clearing before fluorescence imaging. An organic-solvent-based clearing method (uDISCO^50^) was used to clarify the adult mouse brain and zebrafish heart, and a hydrogel-based clearing (CLARITY^2^) method was used to clarify the neonate mouse hearts.

## Acknowledgements

The authors acknowledge the contributions of R. Ardehali and K. Sereti, who assisted with heart sample selection and preparation, as well as R. Kulkarni, H. Chen, and K. Sung, who assisted with organ clearing. The authors also thank X. Wang and Y. Bu for discussions on zebrafish data analysis, thank Z. Yu, and S. Dong for their assistance with instrumentation and GPU-based computation. This research has received funding support from the 1000 Youth Talents Plan of China (P.F.), Fundamental Research Program of Shenzhen (P.F., JCYJ20160429182424047), and the National Heart Lung and Blood Institute (R01HL111437 (T.K.H.), R01HL083015 (T.K.H.), R01HL118650 (T.K.H.), and EB U54 EB0220002 (T.K.H.).

## Author Contributions

P.F. and T.K.H. conceived the research idea and initiated the investigation. J.N., J.L., Y.D., H.Z., P.F., and T.K.H. constructed the system, developed the programs, acquired and processed data, and prepared the manuscript. S.L. and T.S. provided and assisted with HUVEC-HDF cell samples, M.H. provided and assisted with NHBE cell samples, and T.Y. and D.Z. provided and assisted with mouse brains. S.L., T.S., M.H., T.Y., D.Z. and C.H. all advised on image interpretation and manuscript preparation.

## Additional Information

Correspondence and requests for materials should be addressed to (P.F.) or (T.K.H.).

## References

1 Megason, S. G. & Fraser, S. E. Digitizing life at the level of the cell: high-performance laser-scanning microscopy and image analysis for in *toto* imaging of development. Mech. Dev. 120, 1407–1420, doi:10.1016/j.mod.2003.07.005 (2003).

2 Chung, K. et al. Structural and molecular interrogation of intact biological systems. Nature 497, 332–337 (2013).

3 Yang, B. et al. Single-cell phenotyping within transparent intact tissue through whole-body clearing. Cell 158, 945–958 (2014).

4 Ragan, T. et al. Serial two-photon tomography for automated ex vivo mouse brain imaging. Nat. Methods 9, 255–258 (2012).

5 Chalfie, M., Tu, Y., Euskirchen, G., Ward, W. W. & Prasher, D. C. Green fluorescent protein as a marker for gene expression. Science 263, 802–805 (1994).

6 Pawley, J. & Masters, B. R. Handbook of biological confocal microscopy. Opt. Eng. 35, 2765–2766 (1996).

7 Lichtman, J. W. & Conchello, J. A. Fluorescence microscopy. Nat. Methods 2, 910–919 (2005).

8 Power, R. M. & Huisken, J. A guide to light-sheet fluorescence microscopy for multiscale imaging. Nat. Methods 14, 360–373 (2017).

9 Huisken, J., Swoger, J., Del Bene, F., Wittbrodt, J. & Stelzer, E. H. Optical sectioning deep inside live embryos by selective plane illumination microscopy. Science 305, 1007–1009 (2004).

10 Keller, P. J., Schmidt, A. D., Wittbrodt, J. & Stelzer, E. H. Reconstruction of zebrafish early embryonic development by scanned light sheet microscopy. Science 322, 1065–1069 (2008).

11 Keller, P. J. et al. Fast, high-contrast imaging of animal development with scanned light sheet-based structured-illumination microscopy. Nat. Methods 7, 637–642, doi:10.1038/nmeth.1476 (2010).

12 Chen, B. C. et al. Lattice light-sheet microscopy: imaging molecules to embryos at high spatiotemporal resolution. Science 346, 1257998, doi:10.1126/science.1257998 (2014).

13 Ahrens, M. B., Orger, M. B., Robson, D. N., Li, J. M. & Keller, P. J. Whole-brain functional imaging at cellular resolution using light-sheet microscopy. Nat. Methods 10, 413–420, doi:10.1038/nmeth.2434 (2013).

14 Susaki, E. A. et al. Whole-brain imaging with single-cell resolution using chemical cocktails and computational analysis. Cell 157, 726–739, doi:10.1016/j.cell.2014.03.042 (2014).

15 Keller, P. J. & Ahrens, M. B. Visualizing whole-brain activity and development at the single-cell level using light-sheet microscopy. Neuron 85, 462–483, doi:10.1016/j.neuron.2014.12.039 (2015).

16 Vladimirov, N. et al. Light-sheet functional imaging in fictively behaving zebrafish. Nat. Methods 11, 883–884, doi:10.1038/nmeth.3040 (2014).

17 Holekamp, T. F., Turaga, D. & Holy, T. E. Fast three-dimensional fluorescence imaging of activity in neural populations by objective-coupled planar illumination microscopy. Neuron 57, 661–672, doi:10.1016/j.neuron.2008.01.011 (2008).

18 Dodt, H.-U. et al. Ultramicroscopy: three-dimensional visualization of neuronal networks in the whole mouse brain. Nat. Methods 4, 331–336 (2007).

19 Lee, J. et al. 4-Dimensional light-sheet microscopy to elucidate shear stress modulation of cardiac trabeculation. J. Clin. Investig. 126, 1679–1690 (2016).

20 Fei, P. et al. Cardiac light-sheet fluorescent microscopy for multi-scale and rapid imaging of architecture and function. Sci. Rep. 6, 22489 (2016).

21 Guan, Z. et al. Compact plane illumination plugin device to enable light sheet fluorescence imaging of multi-cellular organisms on an inverted wide-field microscope. Biomed. Opt. Express 7, 194–208 (2016).

22 Tomer, R., Ye, L., Hsueh, B. & Deisseroth, K. Advanced CLARITY for rapid and high-resolution imaging of intact tissues. Nat. Protoc. 9, 1682–1697, doi:10.1038/nprot.2014.123 (2014).

23 Ding, Y. et al. Light-sheet fluorescence imaging to localize cardiac lineage and protein distribution. Sci. Rep. 7, 42209, doi:10.1038/srep42209 (2017).

24 Lohmann, A. W., Dorsch, R. G., Mendlovic, D., Ferreira, C. & Zalevsky, Z. Space–bandwidth product of optical signals and systems. J. Opt. Soc. Am. A 13, 470–473 (1996).

25 Brown, M. & Lowe, D. G. Automatic panoramic image stitching using invariant features. Int. J. Comput. Vis. 74, 59–73 (2007).

26 Szeliski, R. Image alignment and stitching: A tutorial. Foundations and Trends^®^ in Computer Graphics and Vision 2, 1–104 (2006).

27 Santi, P. A. et al. Thin-sheet laser imaging microscopy for optical sectioning of thick tissues. BioTechniques 46, 287–294, doi:10.2144/000113087 (2009).

28 Buytaert, J. A. & Dirckx, J. J. Tomographic imaging of macroscopic biomedical objects in high resolution and three dimensions using orthogonal-plane fluorescence optical sectioning. Appl. Opt. 48, 941–948 (2009).

29 Glaser, A. K. et al. Light-sheet microscopy for slide-free non-destructive pathology of large clinical specimens. Nat. Biomed. Eng. 1, 0084 (2017).

30 Maiden, A. M., Rodenburg, J. M. & Humphry, M. J. Optical ptychography: a practical implementation with useful resolution. Opt. Lett. 35, 2585–2587 (2010).

31 Zheng, G., Horstmeyer, R. & Yang, C. Wide-field, high-resolution Fourier ptychographic micxroscopy. Nat. Photonics 7, 739–745, doi:10.1038/nphoton.2013.187 (2013).

32 Hillman, T. R., Gutzler, T., Alexandrov, S. A. & Sampson, D. D. High-resolution, wide-field object reconstruction with synthetic aperture Fourier holographic optical microscopy. Opt. Express 17, 7873–7892 (2009).

33 Fangyen, C. et al. High-speed synthetic aperture microscopy for live cell imaging. Opt. Lett. 36, 148–150 (2011).

34 Gutzler, T., Hillman, T. R., Alexandrov, S. A. & Sampson, D. D. Coherent aperture-synthesis, wide-field, high-resolution holographic microscopy of biological tissue. Opt. Lett. 35, 1136–1138 (2010).

35 Luo, W., Greenbaum, A., Zhang, Y. & Ozcan, A. Synthetic aperture-based on-chip microscopy. Light Sci. Appl. 4, e261, doi:10.1038/lsa.2015.34 (2015).

36 Zheng, G., Lee, S. A., Yang, S. & Yang, C. Sub-pixel resolving optofluidic microscope for on-chip cell imaging. Lab Chip 10, 3125–3129, doi:10.1039/c0lc00213e (2010).

37 Zheng, G., Lee, S. A., Antebi, Y., Elowitz, M. B. & Yang, C. The ePetri dish, an on-chip cell imaging platform based on subpixel perspective sweeping microscopy (SPSM). Proc. Natl. Acad. Sci. U.S.A. 108, 16889–16894 (2011).

38 Luo, W., Zhang, Y., Feizi, A., Göröcs, Z. & Ozcan, A. Pixel super-resolution using wavelength scanning. Light Sci. Appl. 5, e16060 (2016).

39 Xu, W., Jericho, M. H., Meinertzhagen, I. A. & Kreuzer, H. J. Digital in-line holography for biological applications. Proc. Natl. Acad. Sci. U.S.A. 98, 11301–11305, doi:10.1073/pnas.191361398 (2001).

40 Denis, L., Lorenz, D., Thiébaut, E., Fournier, C. & Trede, D. Inline hologram reconstruction with sparsity constraints. Opt. Lett. 34, 3475–3477 (2009).

41 Greenbaum, A. et al. Increased space-bandwidth product in pixel super-resolved lensfree on-chip microscopy. Sci. Rep. 3, 1717, doi:10.1038/srep01717 (2013).

42 Elad, M. & Hel-Or, Y. A fast super-resolution reconstruction algorithm for pure translational motion and common space-invariant blur. IEEE Trans. Image Process. 10, 1187–1193 (2001).

43 Farsiu, S., Robinson, M. D., Elad, M. & Milanfar, P. Fast and robust multiframe super resolution. IEEE Trans. Image Process. 13, 1327–1344 (2004).

44 Vandewalle, P., Süsstrunk, S. & Vetterli, M. A frequency domain approach to registration of aliased images with application to super-resolution. EURASIP J. Adv. Signal Process. 2006, 1–15, doi:10.1155/asp/2006/71459 (2006).

45 Greenbaum, A., Sikora, U. & Ozcan, A. Field-portable wide-field microscopy of dense samples using multi-height pixel super-resolution based lensfree imaging. Lab Chip 12, 1242–1245, doi:10.1039/c2lc21072j (2012).

46 Swoger, J., Verveer, P., Greger, K., Huisken, J. & Stelzer, E. H. Multi-view image fusion improves resolution in three-dimensional microscopy. Opt. Express 15, 8029–8042 (2007).

47 Saalfeld, S. Software for bead-based registration of selective plane illumination microscopy data. Nat. Methods 7, 418–419 (2010).

48 Paxinos, G. & Franklin, K. B. The Mouse Brain in Stereotaxic Coordinates (Gulf Professional Publishing, 2004).

49 Preibisch, S. et al. Efficient Bayesian-based multiview deconvolution. Nat. Methods 11, 645–648, doi:10.1038/nmeth.2929 (2014).

50 Pan, C. et al. Shrinkage-mediated imaging of entire organs and organisms using uDISCO. Nat. Methods 13, 859–867, doi:10.1038/nmeth.3964 (2016).

